# In silico evaluation of the effects of temperature on the affinity of the SV2C ligand UCB-1A to SV2 isoforms

**DOI:** 10.64898/2026.03.19.711868

**Authors:** Rongfeng Zou, Sangram Nag, Vasco Sousa, Anton Forsberg Morén, Miklós Tóth, Yasir Khani Meynaq, Elena Pedergnana, A. Valade, C. Vermeiren, P. Motte, J. Mercier, Xiaoqun Zhang, Per Svenningsson, Christer Halldin, Andrea Varrone, Hans Ågren

**Author notes:** Corresponding author. Rongfeng Zou. Semmelweis University, HUN-REN TKI, Department of Biophysics and Radiation Biology, 1094 Budapest, Hungary.

## Abstract

Synaptic vesicle glycoproteins 2 (SV2) are integral membrane proteins essential for neurotransmitter release and are implicated in neurological disorders including epilepsy and Parkinson’s disease. In the attempt to develop a ligand selective for SV2C, and in collaboration with UCB, UCB-F was identified as a potential candidate. However, the affinity of UCB-F to SV2C was found to be temperature dependent, decreasing by about 10-fold from +4 to 37 degrees. UCB1A was subsequently identified as SV2C ligand displaying in vitro a 100-fold selectivity for SV2C compared with SV2A. In this study we investigated whether the binding of UCB-1A to SV2A and SV2C was affected by the temperature. A combination of experimental binding assay data and molecular dynamics (MD) simulations were used.

The binding studies revealed that UCB1A affinity for SV2A decreased significantly at 37 °C compared with 4 °C, whereas binding to SV2C remained largely unchanged. MD simulations reproduced these observations, namely that ligand RMSD values at 310 K showed that UCB1A binding fluctuated markedly in the SV2A complex, with many trajectories exceeding the 3.0 Å stability cutoff, whereas UCB1A remained relatively well-anchored in SV2C under the same conditions. Structural analysis showed that, while UCB1A adopts a conserved binding pose across all isoforms stabilized by π– π stacking and a hydrogen bond with Asp, SV2C possesses a unique stabilizing feature. In SV2C, Tyr298 is less exposed to the solvent and engages in a persistent hydrogen bond with Asparagine, a structural feature that reinforces pocket stability and limits temperature-induced destabilization. This interaction is absent in SV2A, consistent with its greater temperature sensitivity.

Together, these findings provide a mechanistic explanation for the experimentally observed temperature independence of UCB1A binding to SV2C. More broadly, the results highlight the importance of incorporating physiologically relevant temperatures into SV2 ligand evaluation and demonstrate how combining experiments with simulations can uncover isoform-specific mechanisms of ligand recognition and stability.

## Introduction

Synaptic vesicle glycoproteins 2 (SV2) are integral membrane proteins that play a central role in neurotransmitter release and synaptic function.(2-4) The SV2 family comprises three isoforms (SV2A, SV2B, and SV2C) that are expressed in neurons and endocrine cells with distinct yet overlapping distribution patterns. These proteins contribute to synaptic vesicle priming and calcium-dependent exocytosis, thereby ensuring proper synaptic transmission and homeostasis. Dysregulation of SV2 isoforms has been linked to a variety of neurological and psychiatric conditions, including epilepsy, schizophrenia, and neurodegenerative diseases such as Parkinson’s and Alzheimer’s disease.(5,6) Pharmacological studies have demonstrated that SV2 proteins can be targeted by small molecules to modulate synaptic activity, and clinically used agents such as levetiracetam and brivaracetam have validated the SV2 family as druggable targets.(7) Nevertheless, the molecular mechanisms governing ligand binding and isoform selectivity remain insufficiently characterized, creating challenges for rational therapeutic development.(8) One important complication is the influence of temperature on ligand binding. Biochemical and pharmacological assays are frequently performed under conditions chosen to stabilize proteins rather than to reproduce the physiological environment.(9)

Membrane proteins, transporters, and enzymes in general often show altered binding profiles when measured at low temperature compared with physiological conditions. Low temperatures can stabilize high-affinity conformations by reducing conformational entropy and slowing dissociation, whereas high temperatures increase flexibility and hydration dynamics that can weaken ligand–protein interactions. Despite these well-established principles, temperature effects are rarely considered in early-stage hit validation, which can hinder translation of in vitro findings to therapeutic contexts.(10,11)

For membrane proteins such as SV2, binding assays are frequently performed at 4 °C to slow degradation and conformational changes, thereby extending the stability of the preparation during the experiment. However, ligands can display strong temperature sensitivity in their binding behavior, one example is UCB-F, a ligand displaying high affinity for human recombinant SV2C (pKi=8.4) at 4 °C that decreased by approximately one-fold (pKi=6.8-7.3) at 37 °C (ref.). This temperature-dependent change in affinity for SV2C was consistent with results obtained with in silico modeling (1) and highlights the risk of overestimating binding strength or misinterpreting isoform selectivity when non-physiological conditions are used. In addition, in case of UCB-F, this temperature sensitivity explained the discrepancies between in vitro and in vivo results. UCB-F was developed as ligand for imaging SV2C using positron emission tomography (PET). However, while [^18^F]UCB-F showed in vitro binding to SV2C, no specific binding was observed in vivo in non-human primates (ref).

Therefore, for SV2 proteins in particular, investigation of temperature sensitivity is a key aspect of ligand discovery to reliably assess isoform selectivity of new compounds. The binding to SV2C of a related ligand, UCB1A, is found to be quite temperature insensitive, a property which makes it very useful as a biomarker. Thus, in light of previous findings of temperature dependence we find it imperative to unravel the reason for this circumstance at the molecular level for the particular UCB1A biomarker. In this study, we used a computational approach with MD simulations to study the molecular basis of temperature-dependent binding of UCB-1A to SV2 proteins. The affinity of UCB1A for SV2C was measured at 4 °C and 37 °C.

## Methods

The crystal structure of human SV2A in complex with a small-molecule ligand (PDB ID: 8UO9)(12) was used as the initial template. To construct the SV2A–UCB1A complex, UCB1A was aligned to the co-crystallized ligand in 8UO9. The orientation of each SV2–UCB1A complex within the lipid bilayer was determined using the Orientations of Proteins in Membranes (OPM) database.(14) Complexes were subsequently embedded in a palmitoyl-oleoyl-phosphatidylcholine (POPC) bilayer using the Schrödinger System Builder, with explicit TIP3P water molecules added to fully solvate the system.(15) Counterions (Na^+^, Cl^−^) were included to neutralize the system and approximate the physiological ionic strength. The SV2A–UCB1A model was then subjected to a 10 ns molecular dynamics (MD) relaxation to remove steric clashes and to equilibrate the binding pose. All simulations employed the OPLS4 force field, which provides improved accuracy for protein–ligand interactions in membrane environments.(16)

For SV2C, a homology model was generated using the relaxed SV2A–UCB1A structure as a template.(13) The modeled SV2C–UCB1A complex was refined by a 10 ns MD relaxation to stabilize side-chain conformations and optimize ligand positioning.

To investigate temperature effects, each SV2–UCB1A system was simulated under two conditions: 277 K (4 °C), representing low-temperature assay conditions, and 310 K (37 °C), representing physiological body temperature. Systems were equilibrated with positional restraints prior to production runs. Production simulations were conducted for 100 ns per trajectory, with 10 independent replicas performed for each temperature condition to enhance statistical robustness. Trajectory analyses were performed using a combination of Schrödinger simulation analysis tools and in-house Python scripts.

## Results and discussions

Binding affinities of UCB1A for human recombinant SV2A, and SV2C at 4 °C and 37 °C, were available from in vitro binding studies performed at UCB. At 4 °C, the pKi of UCB1A for SV2A was 7.2 and 6.1 at 37 °C. In contrast, UCB1A binding to SV2C was essentially temperature independent, with pKi values of 8.3 at 4 °C and 8.4 at 37 °C (Table 1). These results demonstrate an isoform-dependent effect of temperature on UCB1A binding: while SV2A interactions are destabilized at physiological temperature, SV2C binding remains stable across the tested conditions.

**Table 1.**
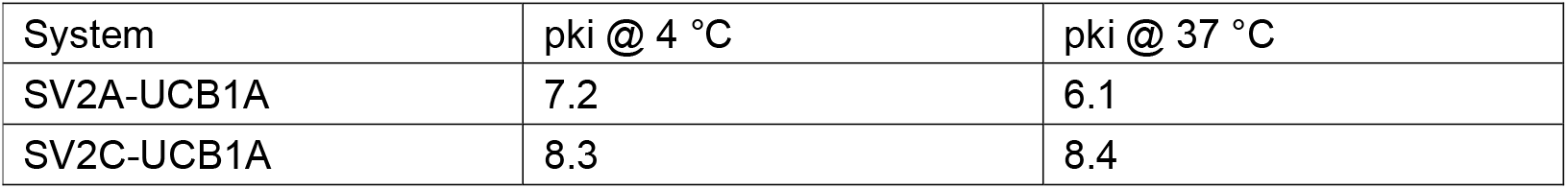
Binding affinities of UCB1A to SV2 Isoforms at different temperature.

| System | pki @ 4 °C | pki @ 37 °C |
| --- | --- | --- |
| SV2A-UCB1A | 7.2 | 6.1 |
| SV2C-UCB1A | 8.3 | 8.4 |

To investigate the molecular basis of the observed phenomenon, we carried out in silico modeling of UCB1A bound to SV2A, SV2B, and SV2C. The binding poses showed highly similar ligand orientations, indicating a conserved binding mode across the three isoforms (Figure 1). In each case, the binding geometry was stabilized by two key interactions: (i) π–π stacking between the aromatic core of UCB1A and a neighboring aromatic residue in the pocket, and (ii) a hydrogen bond formed between the protonated side chain of Asp (modeled as ASH) and a polar substituent of UCB1A, with Asp acting as the hydrogen bond donor. Together, these interactions provide a common structural framework for ligand recognition within the SV2 family.(17,18) Closer inspection of the binding pocket revealed subtle yet meaningful sequence and conformational differences among the isoforms. As highlighted in the upper left corner of Figure 1, the residue at this position varies between isoforms: Gly in SV2A and Asn in SV2C. Such differences are likely to influence the local microenvironment of the binding pocket and, in particular, the dynamics of a nearby Tyr residue close to UCB1A.

**Figure 1.**
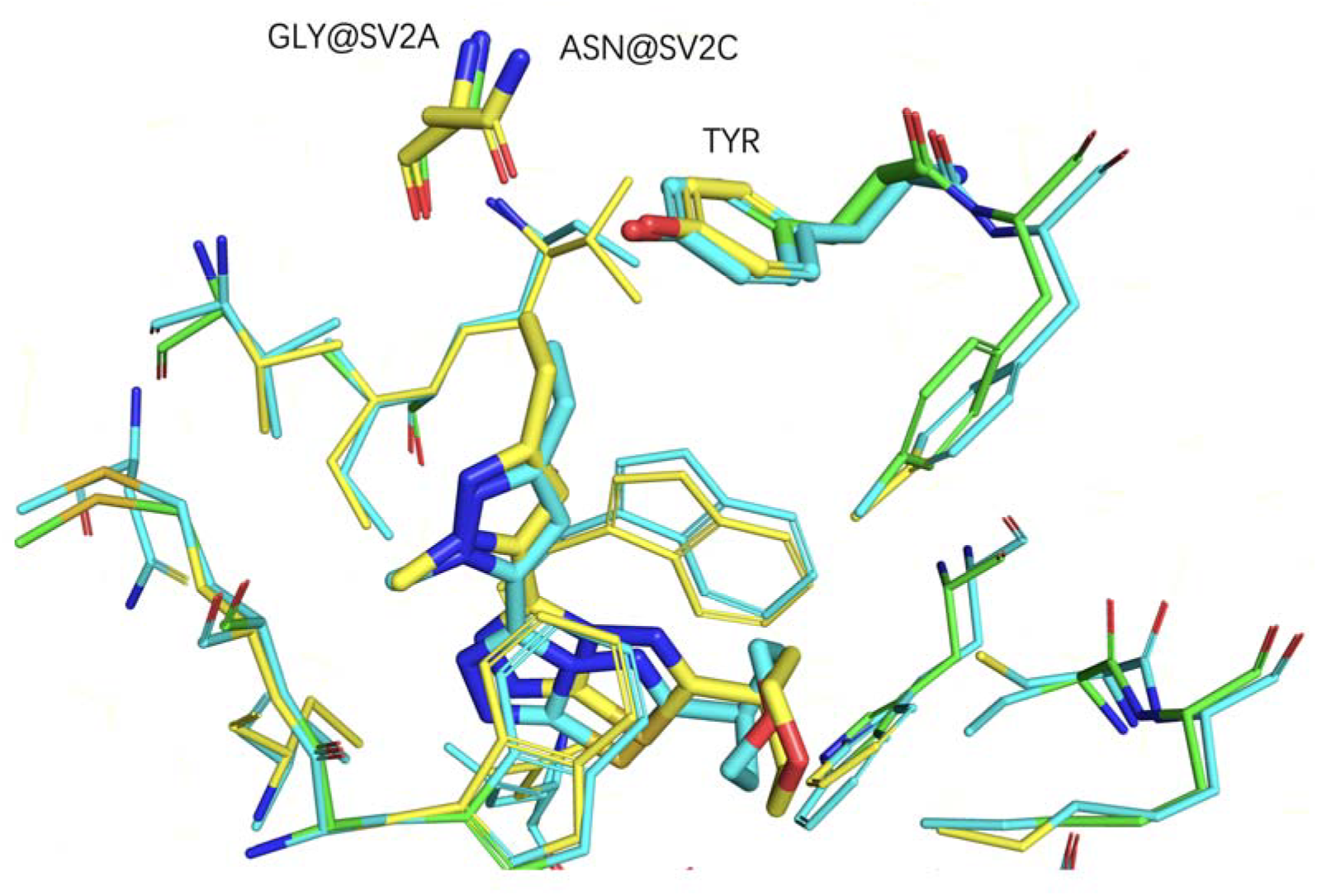
Superimposed binding poses of UCB1A in SV2 isoforms obtained from molecular modelling. Cyan: SV2A–UCB1A; yellow: SV2C–UCB1A.

To investigate the dynamics of these isoform-dependent differences, we performed MD simulations at two temperatures, 277 K (4 °C) and 310 K (37 °C). For each SV2–UCB1A complex, 10 independent replicas were generated at each temperature, providing robust sampling of conformational space and enabling reliable comparison of temperature-dependent effects.

We next analyzed ligand RMSD as a measure of binding stability over the course of the simulations. At 277 K, UCB-1A remained stably bound across all isoforms, with none of the trajectories exceeding the 3.0 Å cutoff. In contrast, simulations at 310 K revealed a marked isoform dependence. For the SV2A–UCB-1A complex, 6 out of 10 replicas exhibited ligand RMSD values greater than 3.0 Å, indicative of partial dissociation or unstable binding. By comparison, only 2 out of 10 trajectories exceeded 3.0 Å in the SV2C–UCB1A complex.

These results demonstrate that increasing temperature destabilizes UCB1A binding to SV2A, whereas SV2C binding is comparatively stable.(19) Notably, the simulation outcomes are in good agreement with the experimental binding assays and results shown in Table 1, which also showed pronounced temperature sensitivity for SV2A but minimal change for SV2C. Together, these findings reinforce the conclusion that SV2C provides a more thermally stable binding environment for UCB1A.

To gain further insight into the structural basis of isoform-dependent binding, we analyzed the solvation environment of the Tyr residue whose hydroxyl group interacts with both UCB1A and neighboring protein residues. The number of water molecules within hydrogen-bonding distance of the Tyr hydroxyl oxygen (O@Tyr) was quantified across trajectories (Figure 2).

**Figure 2.**
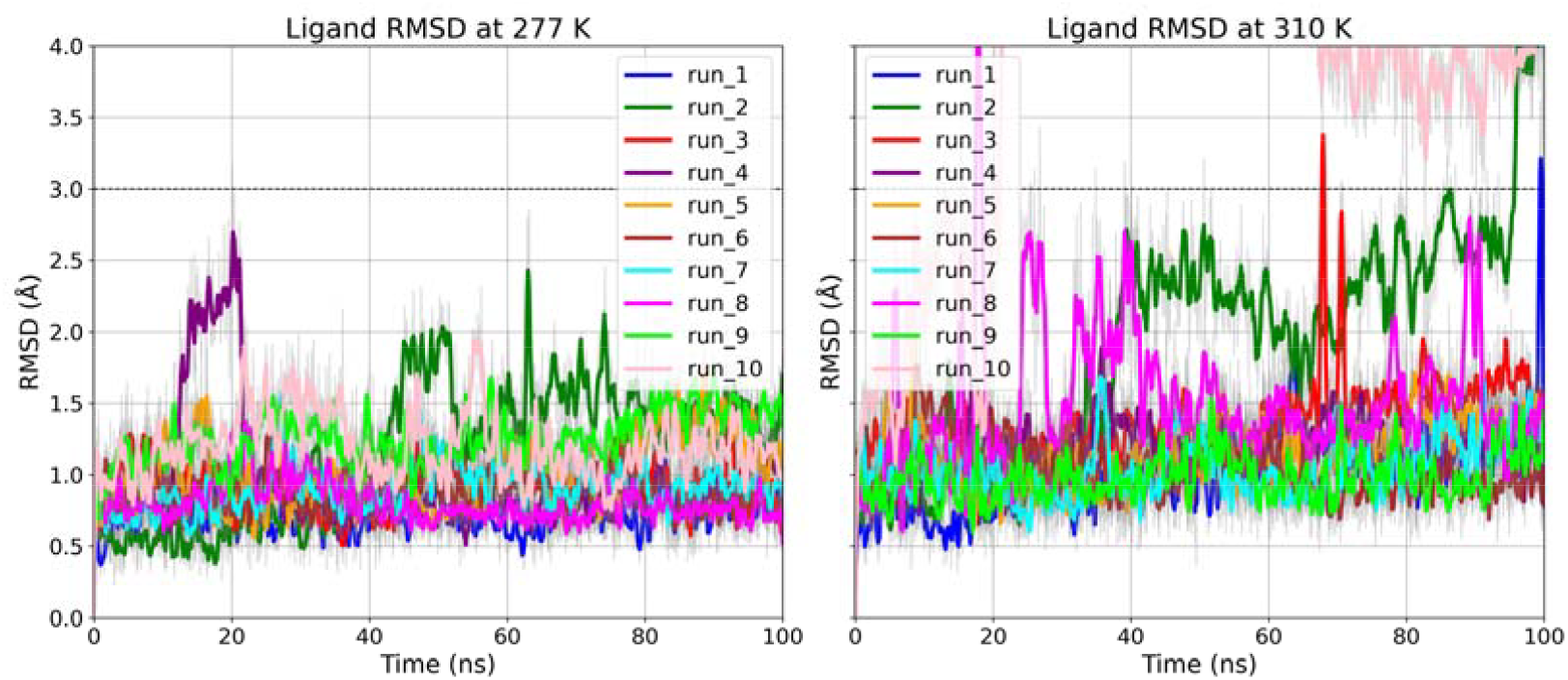
RMSD of UCB1A in SV2A binding pocket at two temperatures. Results from ten independent simulations are shown for 277 K (left) and 310 K (right). The dashed line at 3.0 Å represents the cutoff used to distinguish stable from unstable trajectories.

The results showed that Tyr298 in the SV2C–UCB-1A complex was the least solvated, with an average of ∼1.0 water molecule, compared to higher hydration levels observed in SV2A (1.89). Reduced hydration around Tyr298 in SV2C likely strengthens the hydrogen-bonding network between the Tyr side chain, UCB-1A, and adjacent residues, thereby conferring greater binding stability.(20,21)

Analysis of the SV2C–UCB1A complex revealed a stabilizing hydrogen bond between Tyr298 and Asn, with average donor–acceptor distances of 2.87 Å at 277 K and 3.10 Å at 310 K (Figure 4). Both values are within the range expected for hydrogen bonding, indicating that this interaction is maintained under both temperature conditions. The persistence of this hydrogen bond reduces solvation of Tyr298, thereby strengthening the local interaction network and contributing to the stable binding of UCB1A at physiological temperature. In contrast, this interaction is absent in SV2A due to sequence differences at the corresponding position, consistent with the greater ligand RMSD fluctuations observed in this isoform at 310 K. Taken together, these findings suggest that the Tyr298–Asn hydrogen bond provides a structural basis for the experimentally observed temperature independence of UCB1A binding to SV2C.(22)

**Figure 3.**
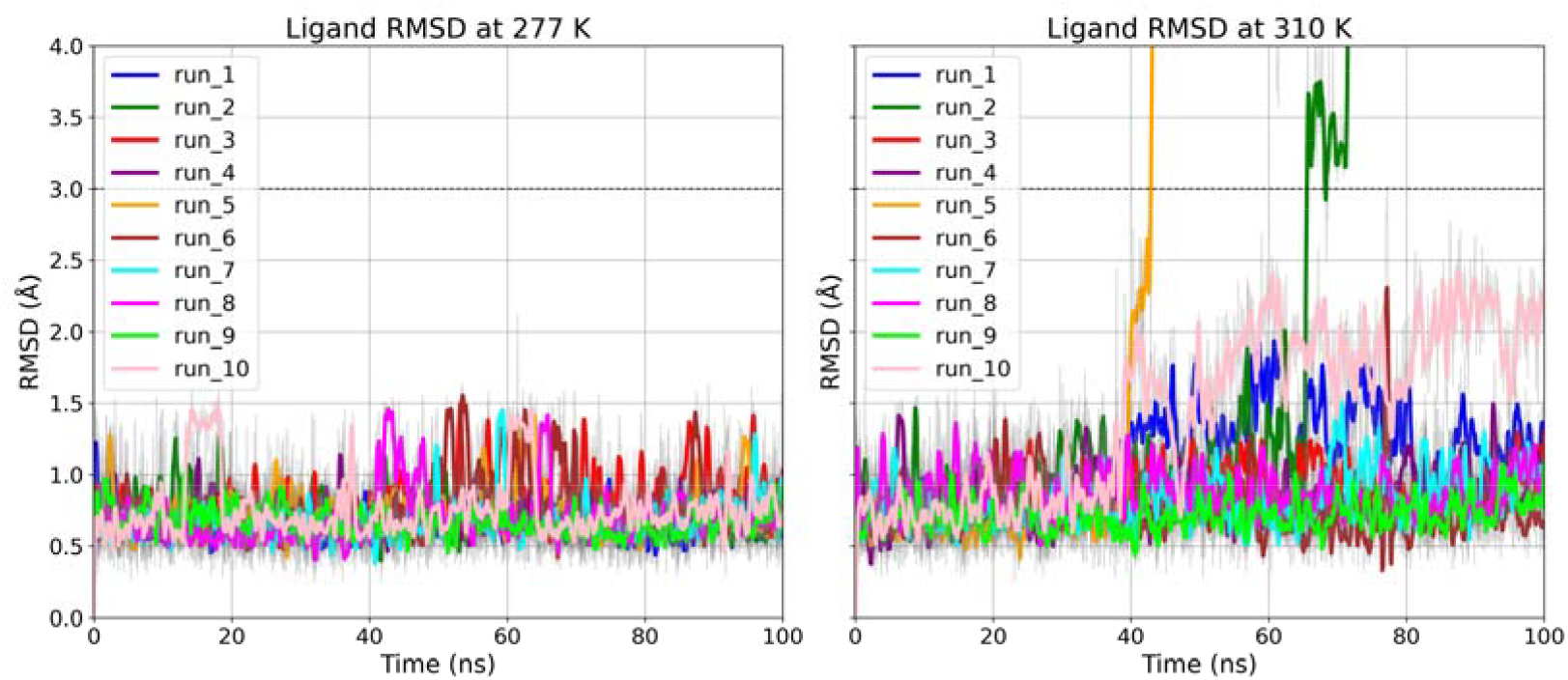
RMSD of UCB-1A in SV2C binding pocket at two temperatures. Results from ten independent simulations are shown for 277 K (left) and 310 K (right). The dashed line at 3.0 Å represents the cutoff used to distinguish stable from unstable trajectories.

**Figure 4.**
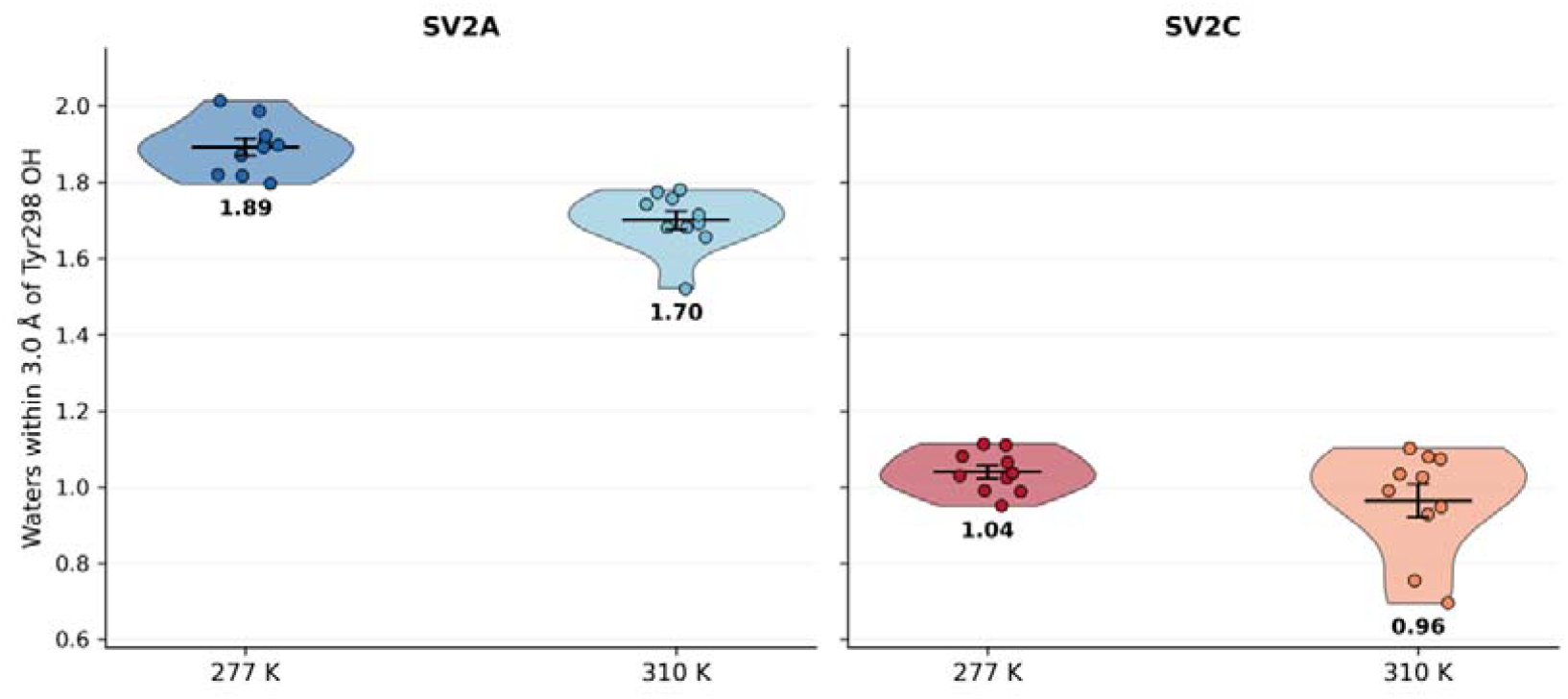
Average number of water molecules within hydrogen-bonding distance of the Tyr hydroxyl oxygen (O@Tyr) across MD trajectories at 277 K and 310 K.

**Figure 5.**
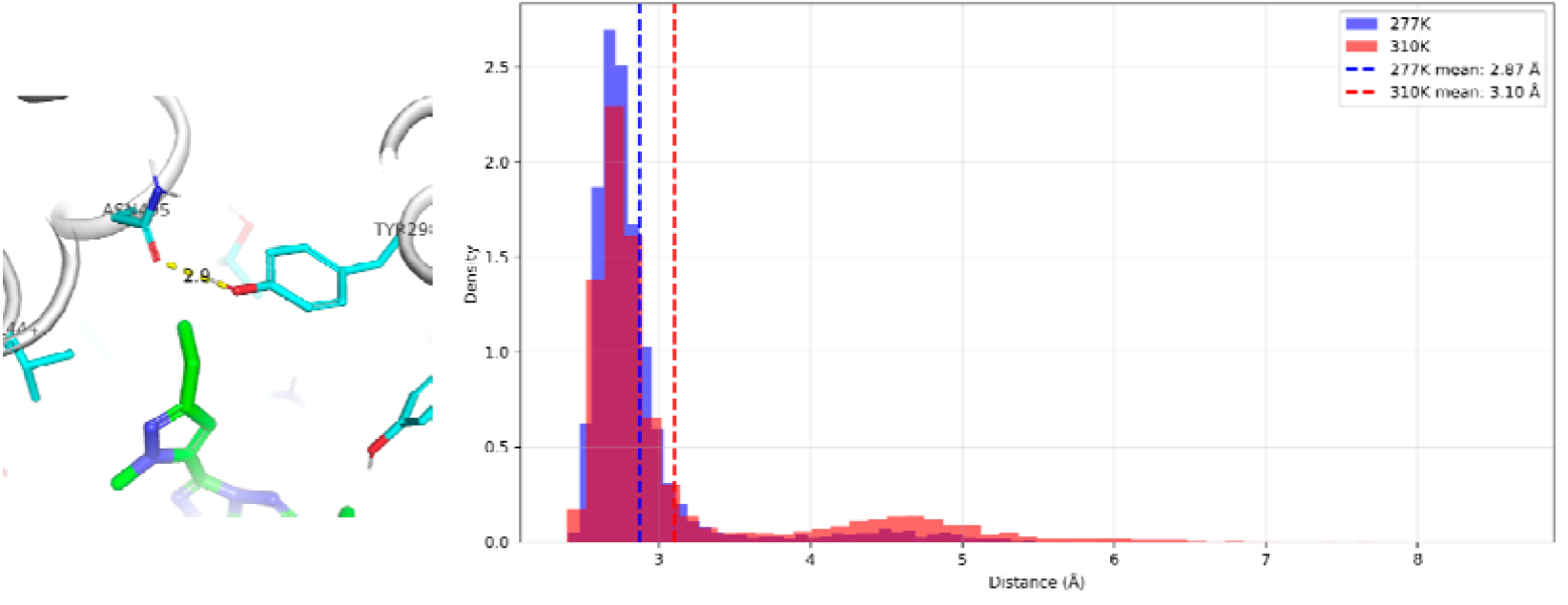
Hydrogen bond analysis of the SV2C–UCB-1A complex. (Left) Representative snapshot showing the hydrogen bond between the side chain of Asp and the hydroxyl group of Tyr298, with UCB1A positioned nearby. (Right) Distribution of Asp–Tyr298 hydrogen bond distances at 277 K (blue) and 310 K (red).

## Conclusions

In this study, we combined experimental binding assays with molecular dynamics simulations to investigate the temperature dependence of UCB1A binding to SV2C. Experimental data showed that UCB1A binding to SV2C remained largely unchanged. Simulations reproduced this trend and evaluated also the effect of temperature on the binding of UCB1A to SV2A. Lligand RMSD analyses showed that a greater number of trajectories exceeded the 3.0 Å stability cutoff in SV2A, while most SV2C trajectories remained below this threshold. Further structural analysis revealed that Tyr298 in SV2C is less solvated than in the other isoforms and engages in a persistent hydrogen bond with Asn, providing a stabilizing interaction absent in SV2A.

Together, these findings highlight an isoform-specific mechanism in which reduced solvation and a stabilizing Tyr298–Asn hydrogen bond buffer SV2C against thermal destabilization. This structural feature explains the experimentally observed temperature independence of UCB1A binding and underscores the importance of considering temperature effects in studies of SV2 ligands.(23,24) Our results demonstrate how integrating biophysical assays with simulations can reveal mechanistic insights into ligand recognition and stability, with ramifications for the rational design of SV2-targeted therapeutics.(25,26)

